# HetMM: A Michaelis-Menten model for non-homogeneous enzyme mixtures

**DOI:** 10.1101/2023.10.10.561792

**Authors:** Jordan Douglas, Charles W. Carter, Peter R. Wills

## Abstract

The Michaelis-Menten model requires its reaction velocities to be measured from a preparation of homogeneous enzymes, with identical or near-identical catalytic activities. However, there are many cases where enzyme preparations do not satisfy this condition, or where one may wish to test the validity of this assumption. We introduce a kinetic model that relaxes this requirement, by assuming there are an unknown number of enzyme species drawn from an unknown probability distribution. This model features one additional parameter over the Michaelis-Menten model, describing the standard deviation of this distribution. We show that the assumption of homogeneity is usually sufficient even in non-homogeneous solutions, and only fails under extreme conditions where Km spans orders of magnitude. We validate this method through simulation studies, demonstrating the method does not overfit to random noise, despite its increase in dimensionality. The two models can be accurately discriminated between even with moderate levels of experimental error. We applied this model to three homogeneous and three heterogeneous biological systems, showing that the standard and heterogeneous models outperform in either case, respectively. Lastly, we show that heterogeneity is not readily distinguished from negatively-cooperative binding under the Hill model. These two fundamentally distinct properties - inequality in catalytic ability and interference between binding sites - give similar Michaelis-Menten curves that are not readily resolved without further experimentation. Our method allows testing for homogeneity and performing parameter inference in a Bayesian framework, and is available online in the user-friendly HetMM package at https://github.com/jordandouglas/HetMM.

## Introduction

One hundred and ten years after its initial publication [1], the equation devised by Leonor Michaelis and Maud Menten is still routinely applied to predicting and explaining biochemical systems, to the point where it is almost synonymous with enzyme kinetics. And like all models, the Michaelis-Menten (MM) model makes several assumptions about the conditions that generated the observed data (see review: [2]). It is assumed that the enzyme concentration is many orders of magnitude less than its substrate, and the latter is assumed to remain stationary for the observed duration of the reaction [3], as does the concentration of the enzyme-substrate complex in the quasi-steady state which characterises the model [4].

Another fundamental assumption, this one so intuitively obvious that it can be easily overlooked, is that the catalysts (and reactants) are homogeneous. That is, the solution is composed of an ensemble of catalysts, each of which displays identical, or near-identical, catalytic activity. However, this condition is often violated in biological systems, even in purified enzyme preparations [5], [6]. In such a scenario, it would be preferable to characterise the kinetics of the catalyst, not as a single entity, but as a *mixture* of catalytic agents, whose catalytic properties are described by some probability distribution.

Indeed, there are numerous cases of non-homogeneous biocatalytic systems, both *in vitro* and *in vivo*. Many genomes express isozymes; enzymes with distinct sequences, but common ancestry and common catalytic activity, such as the multiple aminoacyl-tRNA synthetase gene duplicates that coexist in bacterial cells [7]. In some cases, purified enzyme preparations contain multiple isozymes, such as cytochrome oxidase [6], [8]. But even a single gene can give rise to a heterogeneous mixture of protein products, through transcriptional or translational errors, alternative initiation or splicing, or post-translational modification, among other processes. Otherwise-rare occurrences can be amplified by the environment, for example thermophilic Archaea can adapt to lower temperatures by increasing the rate of methionine-for-leucine mistranslations [9].

Primordial proteins in particular were most likely produced by a low-fidelity and ambiguous genetic code, and hence would have been highly variable [10], [11]. And even proteins with identical sequences, devoid of post-translational modifications, can still fold into a population of distinct structures [12], [13] with a population of catalytic abilities [14], or lack thereof - for instance ion channels [15] and molten globular enzymes [16]. Any of these sources of variability could be further compounded by various degradation effects that may emerge from natural biological processes or from an ill-prepared *in vitro* sample. Catalase, for instance, is known as a “suicide” catalyst because it is degraded by its own substrate [17]. That substrate is hydrogen peroxide, a reactive agent that oxidises proteins; cysteine side chains in particular [18], [19]. Further examples of heterogeneity are illustrated by Brown et al. 2014 [6].

On a coarser scale, heterogeneity may be an intrinsic property of the system that cannot be reduced to some mere idiosyncrasy of a particular gene or protein [20]. This may be the case when studying cells, organelles, or multicellular organisms, in which case the ‘enzyme’ would be a series of complex *in vivo* processes, and the ‘enzyme preparation’ may be living tissue. For instance, Devaux et al. 2023 [21] applied the MM model to quantify oxygen consumption in fish brains.

Generally, heterogeneity is something to avoid in experimental design, but in some cases it may be desirable or even unavoidable. Suppose that not just one, but rather a large sample of enzymes, were to be co-expressed, co-purified, and co-assayed as part of a combinatorial expression library. The assay would measure the kinetic properties of the *population* of forms rather than a single representative from the process that created the enzyme. For instance, experimental investigations into ancestral models typically operate on one enzyme sequence at a time [22]–[25], but this approach could benefit from co-assaying a sample of reconstructions (such as a phylogenetic posterior distribution) to improve the robustness and statistical rigour of the results in a cost-affordable manner.

Some enzymes, such as aspartate kinase [26] and UDP-GlcNAc 2-epimerase [27], display negative cooperativity between binding sites, where binding at one site interferes with the binding affinities at other sites. These interactions are typically modelled using the Hill equation [28]–[30]. However, as discussed by Abeliovich 2005, negative cooperativity is not readily distinguished from a mixture of independent but variable binding sites [31]. They propose the 1/N rule, where a Hill coefficient less than the reciprocal of the number of binding sites likely arose from heterogeneity, rather than negative cooperativity.

Clearly, there are many instances where the standard homogeneous model is unjustifiable, and many more instances where one may wish to test the validity of this assumption. In this article we describe an extension to the Michaelis-Menten model, in which it is assumed that there are an unknown number of catalytic species whose properties are independently drawn from an unknown probability distribution, and thus the enzyme preparation is heterogeneous. We provide a Bayesian model averaging framework, to test whether the standard assumption of homogeneity is adequate, or whether the heterogeneous model is preferred. Lastly, we explore the interactions between the heterogeneous and Hill models, and discuss the limitations of either approach. These methods are implemented in the user-friendly, open-source Heterogeneous Michaelis-Menten (HetMM) package, which employs the BEAST 2 Bayesian inference engine [32]. HetMM is available online at https://github.com/jordandouglas/HetMM.

## Materials and Methods

### Homogeneous and heterogeneous models

Consider the following reaction scheme between substrate A, enzyme E, and product P.

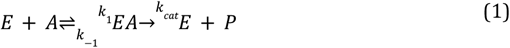

Under the standard (homogeneous) MM model, the expected reaction velocity is equal to

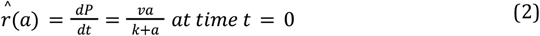

for substrate concentration a, where v is shorthand for V_max_ (the maximum reaction rate) and k for the Michaelis constant K_m_ (the substrate concentration where r is half of V_max_). Under the steady-state approximation, k is defined as:

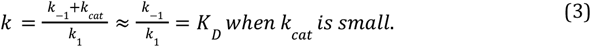

The interpretation of k may change if any of the standard Michaelis-Menten assumptions [2] is not satisfied. Here, we will denote the natural logarithms of v and k by their uppercase symbols V= log v and K = log k. This means that the transformed parameters V and K are not necessarily positive. Now, let us assume that V and K are independently sampled from normal distributions, or equivalently, v and k are independently sampled from log-normal distributions:

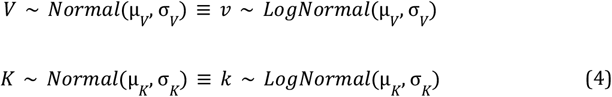

Here, μ and σ are the respective means and standard deviations of V and K. These two normal distributions have probability densities f_V_ and f_K_. Now let us marginalise across all values of K and V according to this distribution:

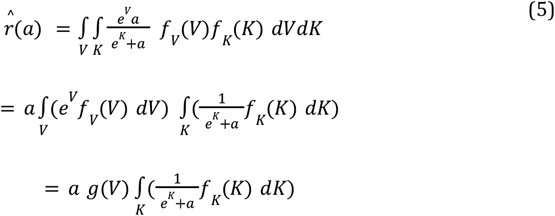

for some function g(V). As this function is independent of a, it is clear that any degree of variation in V cannot be resolved here without further information, such as the relative abundances of the species. Going forward, we will define v_0_ as the mean V_max_ across the species, weighted according to their unknown abundances. However, this simplification is not possible in the case of K [6].

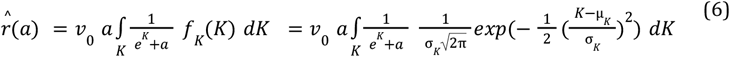

A numerical approximation of this integral is used. This new parameter σK describes the degree of variability in k. As σK approaches zero, the homogeneous and heterogeneous models become indistinguishable, and as σK grows beyond 1-2, the Michaelis constants k span several orders of magnitude (right panel of Fig. 2). As confirmed in Fig. 2, the Michaelis-Menten curve is sensitive to changes in σ_K_ when σ_K_>1. Whereas, when σ_K_<1, the assumption of homogeneity is often adequate, even if untrue.

**Fig 1.**
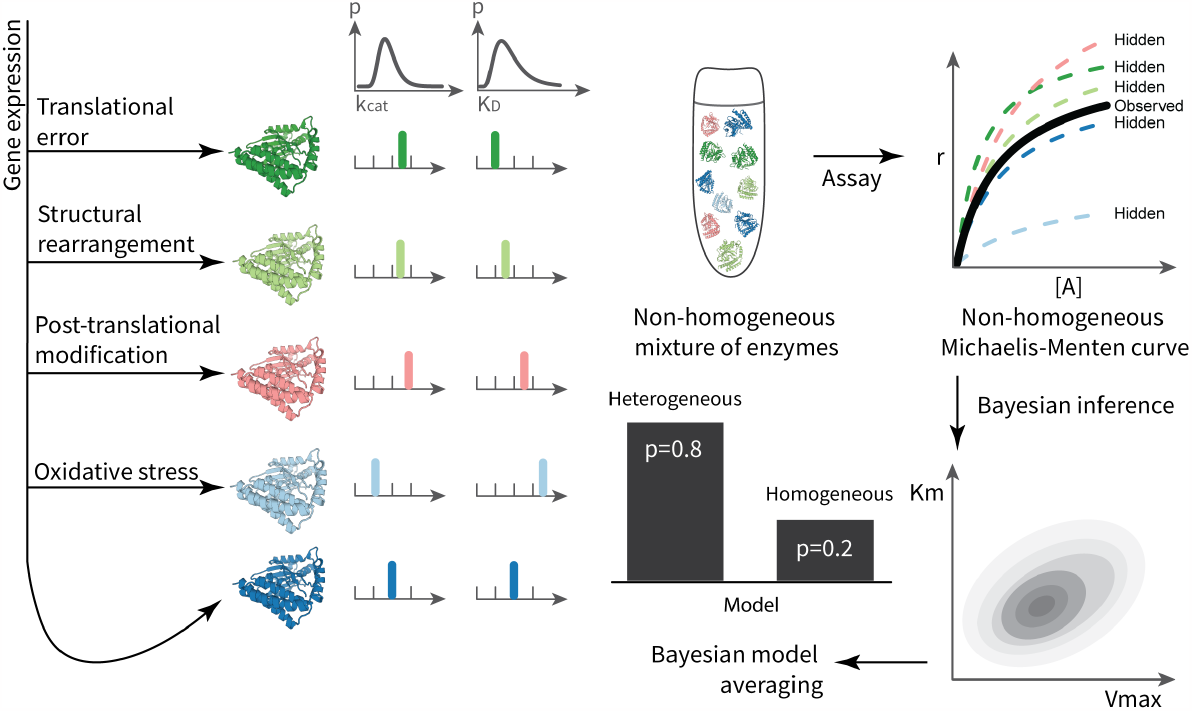
Left: several mechanisms that can lead to enzyme heterogeneity. The properties of each enzyme, kcat and KD, are variable and can be described by some unknown probability distribution, shown at the top. Right: flowchart for testing for homogeneity. The end result is a posterior probability of either model (homogeneous and heterogeneous) being the correct model for the dataset, as well as parameter estimates.

**Fig 2.**
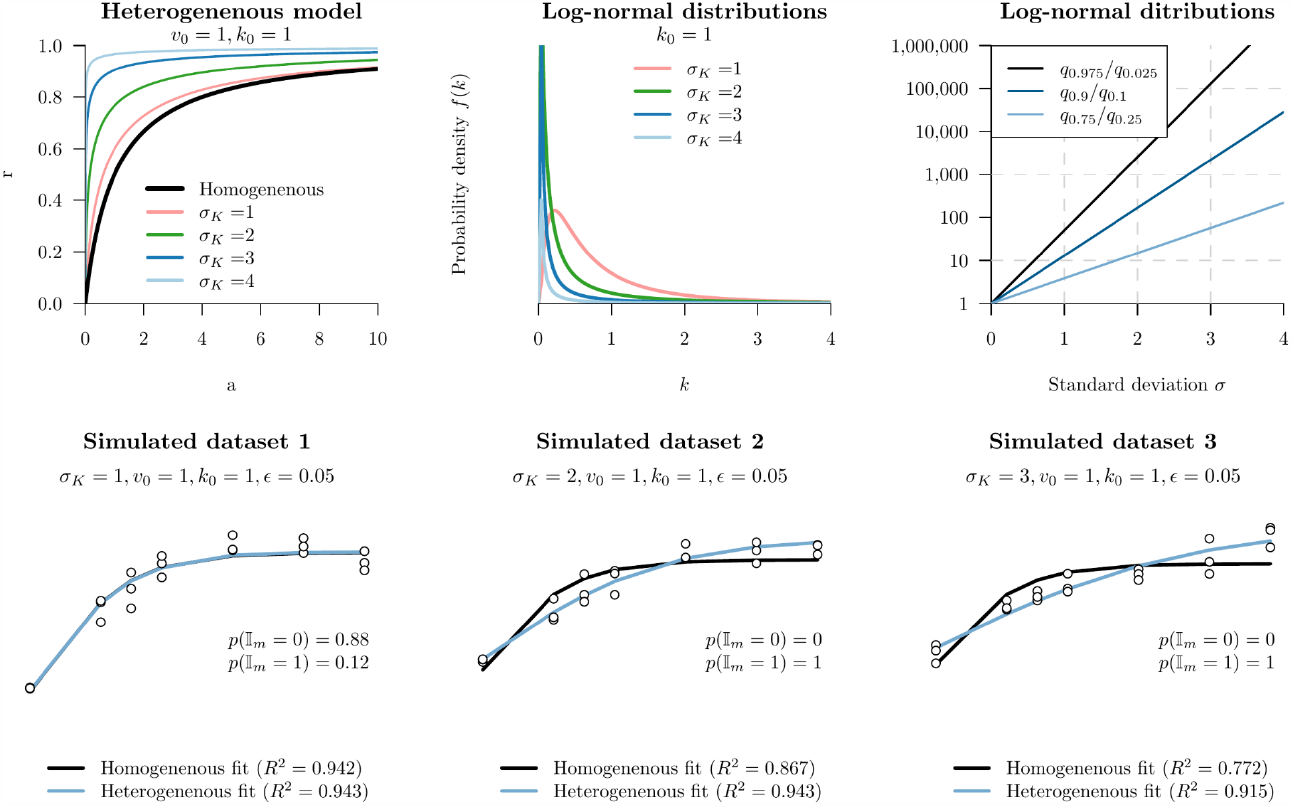
Characterising the effect of σ_K_ on the heterogeneous model. Top row: varying σ_K_ while keeping all other parameters constant changes the Michaelis-Menten curve. This parameter is the standard deviation of a log-normal distribution. The relative difference between upper and lower quantiles q, for varying standard deviations, are shown in the top right figure. Bottom row: three datasets were simulated under the heterogeneous model, with varying σ_K_. The x-axes are concentrations a(log-scale) and the y-axes are reaction velocities r, which were simulated with random error ε. Bayesian inference was run on these datasets, with both models (homogeneous and heterogeneous). In the second two cases (σ_K_ = 2,3), the heterogeneous model is best, because p(I_m_= 1) is large. However, when σ_K_ = 1, the homogeneous and heterogeneous model could not be discriminated between, and therefore the model averaging favoured the simpler model Im= 0.

Overall, this model of heterogeneity assumes there are an unknown number of enzymes whose Michaelis constants fall onto a continuous spectrum of binding affinities and catalytic rates. In practice, the various k values of the mixture need not be log-normally distributed, however this flexible distribution is significantly more relaxed than the assumption of all catalytic species having identical k. This continuous model differs from the heterogeneity model by Brown et al. 2014 [6], which assumes a small number of individually homogeneous species (e.g., two species) that are weighted according to their relative abundances.

### How heterogeneous is heterogeneous?

Before the distinction between homogeneity and heterogeneity makes any sense, we must first quantify how disparate the catalytic species should be so that their population is observably non-homogeneous. Suppose, for instance, that an ensemble of enzymes were composed of several species, whose K_m_ ranged from 1 to 1.1 units (if the species were to be characterised in isolation). In this case, the system would be effectively homogeneous. As demonstrated by Brown et al. 2014, the assumption of homogeneity is likely sufficient for the case of two species, provided that the ratio between their K_m_ is no more than 20 [6].

Moreover, as shown in Fig. 2 (bottom right), the standard deviation σ_K_ should be > 1 in order for the relative difference between the upper and lower extremes of k (among the population of enzymes) to be multiple orders of magnitude apart, and for the two models to have distinguishable properties. This logic forms the basis of our prior distribution for σ_K_, which is shown in Table 1 along with the other priors.

**Table 1.** Prior distributions used in this study. Note that a Log-normal distribution’s μ and σ are the mean and standard deviation in log-space. In a real world example, the priors for v0 and k0 should be adjusted to reflect the units of measurement and other a priori information about the system being studied.

| Parameter | Prior | Notes |
| --- | --- | --- |
| *General case* |  |  |
| $v_0$ | Log-normal($\mu = 1, \sigma = 2$) | Uninformative prior |
| $k_0$ | Log-normal($\mu = 1, \sigma = 2$) | Uninformative prior |
| $\sigma_K$ | Log-normal($\mu = 1, \sigma = 0.5$) | Ensures the heterogeneous model is "heterogeneous" |
| $\epsilon$ | Log-normal($\mu = -1.5, \sigma = 1$) | A reasonable level of experimental error |
| $I_m$ | Uniform($\{0, 1\}$) | Both models are equal *a priori* |
| *Hill model* |  |  |
| $h_0$ | Beta($\alpha = 2, \beta = 4$) | Shifts $h$ away from 1 |
| $I_h$ | Uniform($\{-1, 0, 0, 1\}$) | Hill and not-Hill are equal *a priori* |
| *Simulation studies* |  |  |
| $k_0$ | Log-normal($\mu = 1, \sigma = 0.5$) | |
| $v_0$ | Log-normal($\mu = 1, \sigma = 0.5$) | |
| *Hb dataset* |  |  |
| $v_0$ | 100 | Fixed at 100% saturation |
| *MnP dataset* |  |  |
| $I_h$ | 0 | Hill model disabled (only one active site) |
| *CFTR dataset* |  |  |
| $h_0$ | Beta($\alpha = 30, \beta = 10$) | Low probability of $h < 0.5$ or $h > 2$ (two active sites) |

### Bayesian inference and model averaging

Parameter estimation is performed using Bayesian inference, allowing for robust estimation of parameters and their credible intervals, as well as the model indicator I_m_ which governs the use of the homogeneous or heterogeneous MM model (i.e., model averaging). Let *D* be the data measured from a kinetic assay, consisting of a vector of substrate concentrations a = a_1_, a_2_, …, an and empirical reaction rates r = r_1_, r_2_, …, rn. Given model parameters θ, the likelihood is calculated by comparing observations with their expected values, conditional on substrate concentration a, using the equation

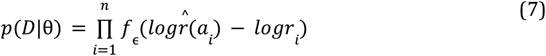

where the experimental errors between expected and observed values are assumed to follow a normal distribution, in log-space. This normal distribution, with probability density fε has mean 0 and standard deviation ε. Modelling experimental error in log-space accounts for the fact that reaction rates are non-negative, while also accommodating for the common scenario where the magnitude of error increases with substrate concentration, i.e., heteroscedasticity.

This model consists of five parameters θ = (v_0_, k_0_, σ_K_, ε, I_m_). If I_m_=0, then the enzymes are assumed to be homogeneous, in which case reaction velocities are predicted using Eqn. 2, where v=v_0_ and k=k_0_. σ_K_ is not being used in the model. Whereas, if Im=1,, then the enzymes are assumed to be heterogeneous, in which case reaction velocities are calculated using Eqn. 6, where μ_K_ = log k_0_ - 0.5σ 2 so that μK reflects the mean of k in real-space, rather than log-space. In this model, all five model parameters in θ are being used.

Lastly, the posterior density is

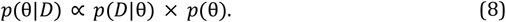

Our prior distributions p(θ) are summarised in Table 1. The posterior distribution is sampled using Markov chain Monte Carlo (MCMC), as implemented in BEAST 2 [32]. Although BEAST 2 was designed for phylogenetic inference, its use of efficient proposal kernels, including Bactrian [33], adaptable variance multivariate normal [34], and adaptable operator sampler [35] kernels, makes it an attractive engine for the purposes of this model. MCMC chains are run until the effective sample sizes of all parameters exceed 200, as diagnosed by Tracer [36].

Lastly, we wish to address a point of potential confusion concerning the varying use of log-normal distributions. We have used log-normal distributions in three different contexts here. First, in Eqn. 4, we assume that the properties of enzymes in the population are distributed in a log-normal fashion, i.e., log-normal(μ_K_, σ_K_). Second, in Eqn. 7, we assume the experimental error is drawn from log-normal(0, ε) - this distribution describes random error in the experimentation process. Lastly, in Table 1, we use the log-normal as prior distributions - these distributions describe information about any *a priori* expectations of a parameter, before performing any analyses. As shown in Fig. 2, log-normal distributions are flexible, they have positive domains, and their shapes are determined by only one parameter, σ, making them readily interpretable.

### Hill model extension

We describe a heterogeneous extension to the Hill model. The Hill equation is an empirical model that allows interaction between binding sites or the compartments of a tissue, e.g., muscle [28]–[30]. The nature and degree of this interaction is described by the Hill coefficient h>0:

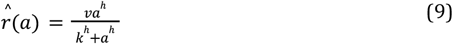

When h>1, this indicates positive cooperativity between binding sites (giving a sigmoidal MM curve), and h<1 indicates negative cooperativity (giving an MM curve with a sharpened hyperbola). When h=1, there is no interaction, and the Hill model is equivalent to MM. Many have argued that the Hill model is merely a descriptive model, and lacks a strong theoretical basis [30]. In any case, like the MM model, the Hill model assumes homogeneity. Thus, under the heterogeneous Hill model:

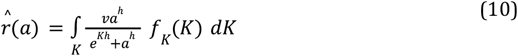

The posterior distribution of this model θ_h_ is estimated using MCMC from the posterior density (Eqn. 8). However, there are two additional parameters in θ_h_ that are absent from the heterogeneous MM model. These two parameters govern whether the Hill coefficient is positive, negative, or neutral. h is equal to h0 when I_h_=-1, 1 when I_h=0_, and 1/_h_0 when I_h_=1, where 0 < _h_0 < 1 is the Hill coefficient in the case of negative cooperativity, and its reciprocal for positive cooperativity. This parameterisation ensures symmetry in their prior distributions, and places the probability mass away from h=1 so that the three models are distinct. Moreover, their prior distributions should reflect prior knowledge about the number of binding sites, and place the majority (or all) of their probability densities within the range (1/N, 1) and (1,N), respectively, where N is the number of binding sites in each enzyme. Inferring the value of the model indicator Ih during MCMC can be used to test whether the Hill model is applicable to a given dataset, and if so, then the direction in which cooperativity acts.

## Results

### Validation on simulated data: the model is well-calibrated

In order to test how accurately our method can recover parameters from datasets generated by the same model, we simulated two hundred datasets and recovered the parameters on each of them using Bayesian MCMC. These datasets were simulated using parameters that were randomly sampled from the prior distribution. This experiment was designed to test a) whether the true parameters can be recovered, and b) whether the true model (homogeneous or heterogeneous) can be recovered. It would be undesirable if, for instance, the heterogeneous model was disproportionately selected even on homogeneous data. The conditions of these simulations mirrored a typical experimental setup, with three sets of reaction rates independently measured per substrate concentration, across seven different concentrations. These seven concentrations flanked the Michaelis constant *k*, which was sampled from the prior with a mean value of 3 units. These concentrations were 0.5, 2.5, 5, 10, 50, 250, and 1000 units.

These results provide confidence that the method is able to recover the true parameters and model (Fig. 3). In the case of continuous parameters, the true value was in the 95% credible interval approximately 95% of the time. And in the case of the model indicator I_m_, the true model was identified, with over 95% support, the majority of the time. In some cases, the model could not be confidently resolved, where 0.05 < p(I_m_ = 1) < 0.95. These simulation studies provided no evidence of systematic “overfitting”, where the model with a larger dimensionality (i.e., the heterogeneous model) was favoured because it captures random noise. Moreover, discrimination between the two models can be achieved even on datasets with moderate degrees of experimental error, where ε is less than around 0.5. The model is therefore quite robust to random error - a standard deviation of ε=0.5 means that 95% of the measured reaction velocities at a given concentration can span up to 10 fold, and yet the true model (homogeneous/heterogeneous) can often still be recovered. We performed the same experiment for the homogeneous Hill model and found similar results (Supporting information). This stringent validation method (simulation-based calibration [37]) is becoming increasingly performed when testing Bayesian models in biological settings [38]–[40].

**Fig 3.**
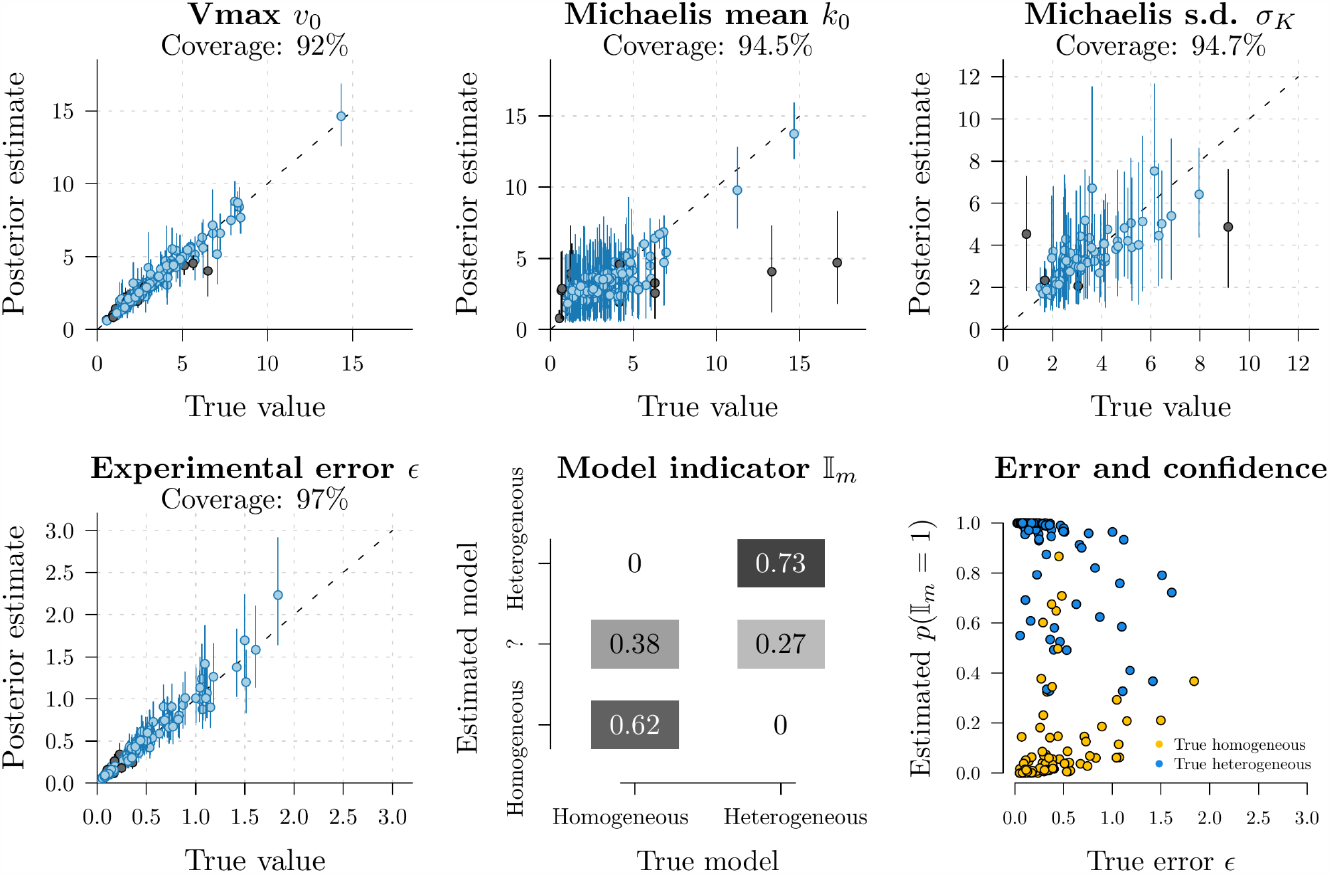
Well-calibrated simulation study. Two hundred datasets were simulated, and the parameters used to generate them were recovered with Bayesian MCMC. The coverage is the percentage of simulations where the true parameter is in the 95% credible interval (blue lines), compared to the times when it is wrong (black lines). These results show that coverage is close to 95%, thus providing confidence in the ability of the method to recover parameters. σ_K_ is not part of the homogeneous model, and therefore its coverage is only calculated when p(I_m_ = 1) > 0.95. In the bottom middle panel, the estimated model is indicated if it has more than 0.95 posterior support, while ‘?’ denotes uncertainty (i.e., 0.05 < p(I_m_ = 1) < 0.95).

We also explored the possibility of resolving the newly introduced heterogeneous model from the standard Hill model. To do this, we simulated and performed Bayesian inference on a further 400 datasets, but this time they were simulated under 6 different models: homogeneous and heterogeneous, crossed with positive, negative, and neutral Hill. Half of these datasets were simulated at 7 substrate concentrations, and the other half with 14 concentrations (3 replicates per concentration). These results suggest that the Hill model, which is often interpreted as describing interactions between binding sites, is in many cases, indistinguishable from the heterogeneous model, i.e., the models are non-identifiable (Fig. 4). Doubling the number of observations improved the resolution but did not solve the issue. In simple terms, a non-homogeneous enzyme preparation may give the appearance of an interaction between binding sites, and a system whose binding sites are interacting may give the appearance of an inequality in their catalytic abilities. These two distinct mechanisms are not readily disentangled without further information, or a large volume of high quality kinetic data.

**Fig 4.**
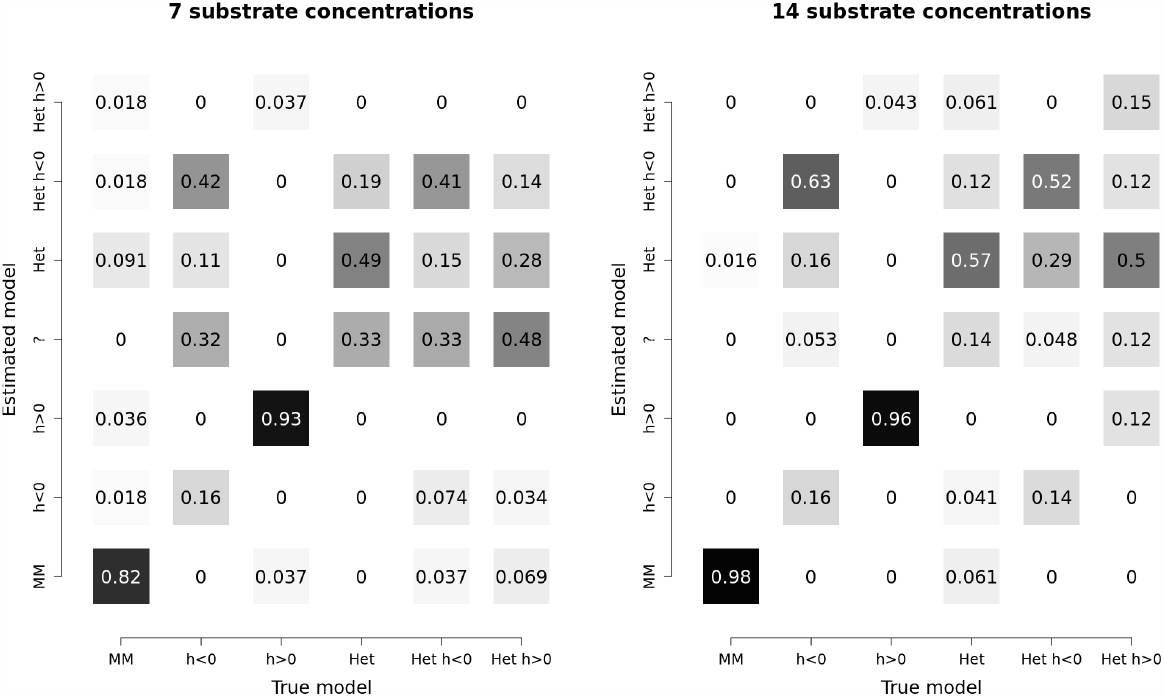
Datasets were simulated under six models: homogeneous Michaelis-Menten (MM), homogeneous negative Hill (h<1), homogeneous positive Hill (h>1), heterogeneous Michaelis-Menten (Het), heterogeneous negative Hill (Het h<1), and heterogeneous positive Hill (Het h>1). Bayesian inference was performed on each dataset to estimate the model and its probability. When any model is identified with greater than 50% posterior support, it is indicated accordingly on the y-axis, and when no single model can be identified, it is labelled with a ‘?’. These datasets were simulated with three replicates of seven (left) and fourteen (right) concentrations of substrate, using parameters sampled from the prior. These results suggest that the Hill model and heterogeneous models are often non-identifiable and the correct model cannot always be recovered, even on larger datasets, corroborating the findings of Abeliovich 2005 [31].

### Negative control: homogeneous biological systems

We tested our method on three kinetic datasets that were obtained from putatively homogeneous enzyme preparations. In each case, we tested for both homogeneity and use of the Hill coefficient in a joint Bayesian analysis.

First, we considered the original experiments carried out by Michaelis and Menten in 1913 [1], which have since been translated into English [41]. They measured the hydrolysis of sucrose (into glucose and fructose) by an enzyme then-known as invertase (EC. 3.2.1.26), across seven sucrose concentrations. Second, we considered the hydrolysis of o-nitrophenyl-β-D-galactopyranoside (ONPG) by β-galactosidase (EC. 3.2.1.23). These data come from an example used in the renz package for R [42], describing eight experimental replicates, each one carried out by a different group of second-year college students. Third, we considered the human haemoglobin (Hb) dataset from Severinghaus 1979 [43], which has been described as a “gold standard” for cooperative binding data [44]. The Hill model is often applied to modelling Hb, as it describes the interaction between its binding sites. These three datasets come with varying standards of experimental precision, as quantified by ε (Table 2). The Michaelis-Menten dataset had the lowest experimental error (ε=0.033), while unsurprisingly, ONPG, which was aggregated from eight student groups, was comparatively noisy (ε=0.35).

**Table 2.**
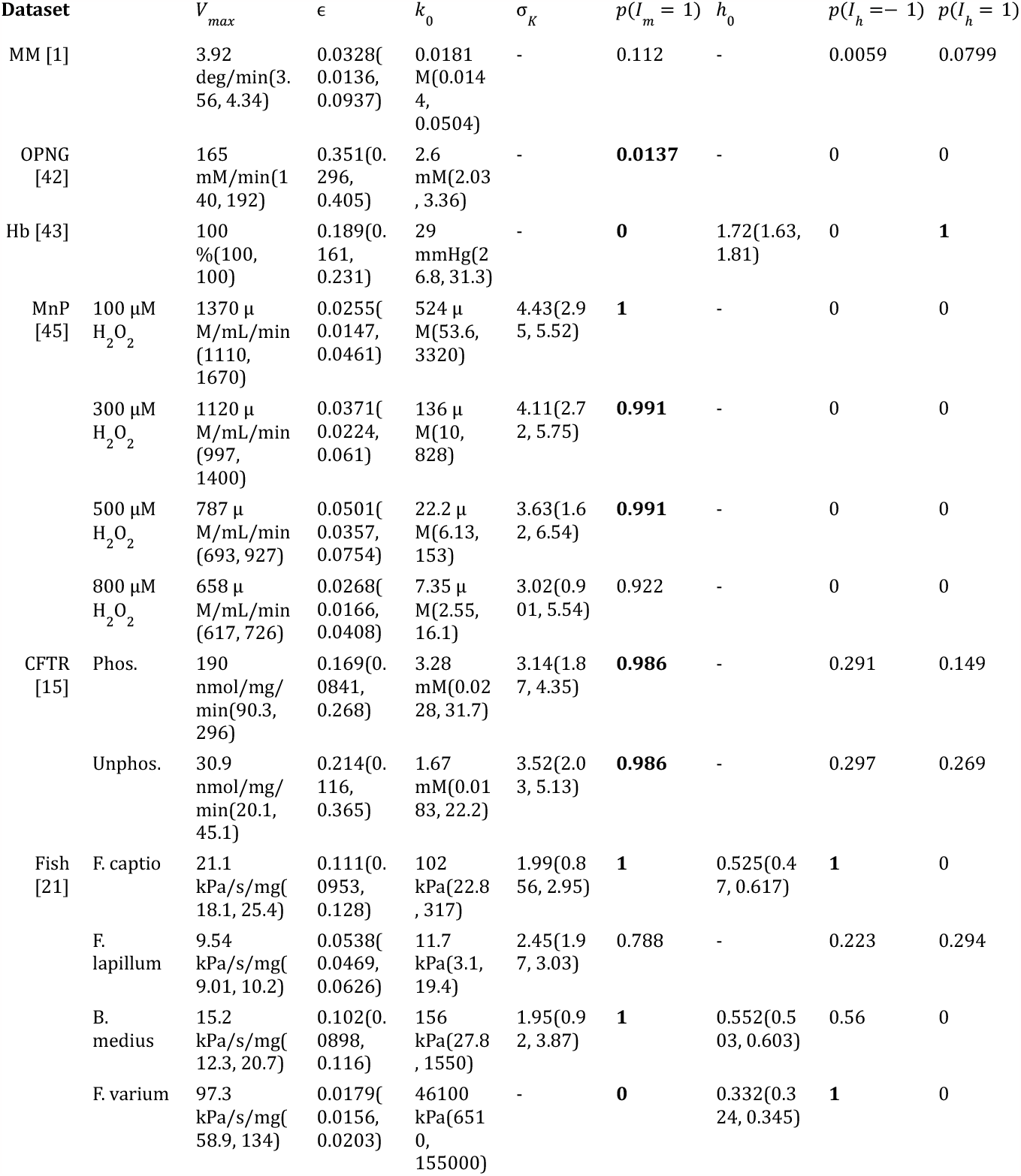
Testing biological datasets for homogeneity and their use of the Hill coefficient. Shown are median parameter estimates (and 95% credible intervals), rounded to 3 significant figures. The estimates for σ_K_ and h0 are omitted when the heterogeneous and Hill models are not being used, respectively.

In all three cases, the homogeneous model was selected (Table 2 and Fig. 5). Specifically, the OPNG and Hb datasets strongly rejected the heterogeneous model, with p(I_m_=1)<0.05, while MM was less confident. The Hill model was rejected by MM and OPNG, and selected for Hb, with h=1.72, consistent with positive cooperativity between binding sites. These experiments further corroborate our simulation studies, confirming that our model, and its extra parameter, do not have a tendency to overfit to random noise in the data.

**Fig 5.**
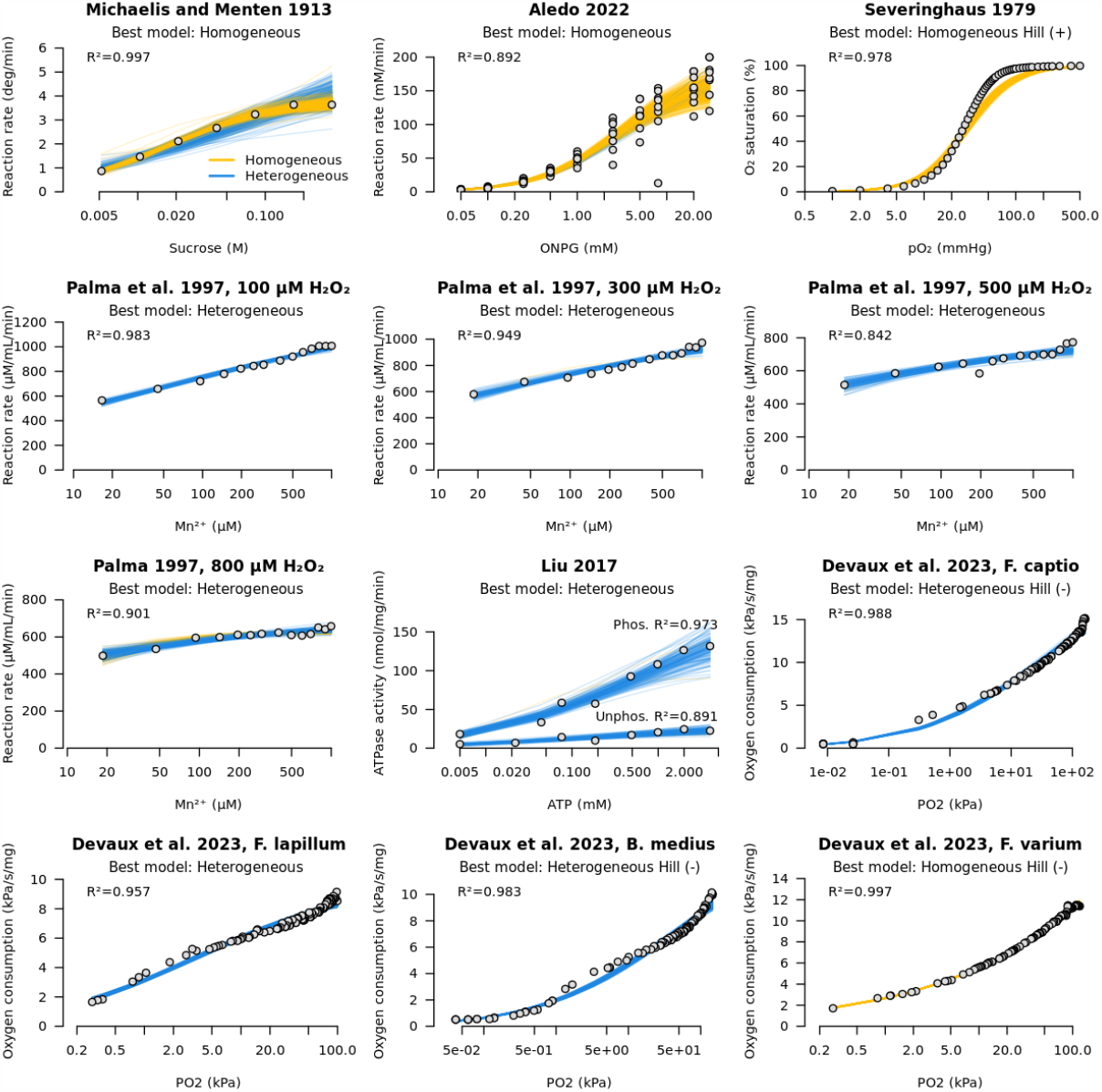
Testing biological datasets for homogeneity and their use of the Hill coefficient. Observations are denoted by open circles, and each coloured line represents a sampled fit, under the posterior distribution. Yellow lines indicate a homogeneous-model-sampled fit, and blue for heterogeneous. Note that the x-axes are on a logarithmic scale.

### Positive control: heterogeneous biological systems

Finally, we tested our method on three biological systems, each suspected to be non-homogeneous. These datasets were tested for both homogeneity and use of the Hill coefficient. Where appropriate, the Hill coefficient was heavily constrained *a priori* within (1/N, N) for N binding sites, in accordance with the 1/N rule [31].

First, we considered an enzyme that is degraded by its own substrate. This enzyme is manganese peroxidase (MnP; EC. 1.11.1.13), and it catalyses the oxidation of Mn^2+^ to Mn^3+^ in the presence of H_2_O_2_. However, H_2_O_2_ is a reactive oxygen species, which decomposes into an even more reactive agent HO- in the presence of metal ions, leading to protein degradation [18]. Palma et al. 1997 showed that Mn^2+^ exerted a positive effect on enzymatic activity, whereas excess levels of H_2_O_2_ in fact lowered the reaction rate [45]. We applied our method to initial oxidation rates at varying levels of Mn^2+^, which were initiated at four levels of H_2_O_2_. When testing for homogeneity, our method rejected the assumption on all four datasets, with p(I_m_=1) ranging from 0.92 to 1.0. These results are consistent with reactive products of hydrogen peroxide attacking the protein ensemble, leading to subpopulations of enzymes with severely impaired catalytic activities, while still remaining functional. MnP is a monomer with a single active site [46], and therefore its Hill coefficient was fixed at 1.

Variability in Michaelis constant is the best explanation for the non-MM behaviour in this case.

Second, we considered a transmembrane ion channel that adopts “open” and “closed” conformations. This channel is the human cystic fibrosis transmembrane conductance regulator (CFTR) protein (EC. 5.6.1.6), and it binds ATP in order to open the channel, enabling the flow of chloride-ions along their electrochemical gradient [47], [48]. The channel closes upon ATP hydrolysis. The catalytically active (open) state is promoted by phosphorylation of the R domain [15]. As this is a complex multi-state system, it is reasonable to hypothesise that the active sites may take on a broad distribution of catalytic affinities, and therefore behave heterogeneously. CFTR contains two ATP binding sites [49], and therefore its Hill coefficient should be no less than 0.5 and no more than 2 in order to explain cooperative binding. This constraint was reflected in our prior distributions.

Indeed, our method favoured the heterogeneous model, with h=1, in both datasets by Liu et al. 2017 (phosphorylated and dephosphorylated CFTR). The Hill model was not selected and therefore heterogeneity is a much better explanation for the non-MM behaviour displayed by CFTR than cooperative binding. Without invoking a multistate model of CFTR domain rearrangement [15], this coarse-grained description of heterogeneity remains the best explanation.

Third, we considered an *in vivo* assay on fish brains. Devaux et al. 2023 evaluated oxygen consumption in the brains of four species of triplefin fish [21]. The efficiency of the electron transfer chain is likely to vary from mitochondrion-to-mitochondrion and cell-to-cell, and therefore a living tissue like this is likely to be intrinsically non-homogeneous. In contrast to our previous systems, constraining the Hill coefficient to reflect the number of binding sites is no longer straightforward as this is a living tissue and cooperativity could come in many forms. Across the four fish brains, our method selected either the heterogeneous model, the Hill model with negative cooperativity, or both. Thus, it is unclear which explanation - heterogeneity or negative cooperativity - is most suitable without further information.

## Discussion

In this article we presented a model that tests for and captures heterogeneity inherent in the Michaelis constants (Km) of enzymic systems. The method was validated using simulated data (Fig. 3) and then applied to three biological datasets where it greatly outperformed the standard Michaelis-Menten model (Fig. 5). This model features an additional parameter σ_K_ that describes the standard deviation of Michaelis constants between active sites, which are assumed to be independently drawn from a log-normal distribution. But despite the additional dimensionality, there is no evidence of this model systematically overfitting to random noise in the data. Moreover, the method is quite tolerant to random experimental error. The true model can often be recovered even when the observed reaction velocities span up to one order of magnitude across replicates, but of course less error is always preferred (Fig. 3). Corroborating the results of Brown et al. [6], we showed that the standard assumption of homogeneity imposed by the Michaelis-Menten model is likely to be sufficient in most cases, and failing only under extreme conditions when the different enzymes’ Michaelis constants span orders of magnitude (Fig. 2). Under these conditions, some enzymes must be significantly less proficient than others, but still active, otherwise their catalytic activities would not be detected.

Heterogeneity is often indistinguishable from negative-cooperation under the Hill model [31], which has been widely criticised for lacking a strong theoretical basis [30]. There exists a twilight zone, in which independently heterogeneous enzymes give the appearance of being non-independently interfering, and vice versa, and the two fundamentally distinct properties give rise to similar hyper-rectangular Michaelis-Menten curves. The two properties are not readily disentangled without further information, such as the number of binding sites per enzyme [31]. This represents a fundamental limitation in both models, as well as the broader approach of inferring the behaviour of molecular processes from two dimensional hyperbolic curves.

We tested six biological systems for their homogeneities and cooperative binding. These systems were a) sucrose hydrolysis from Michaelis and Menten 1913 [1], b) ONPG hydrolysis from second-year college students [42], c) oxygen binding by haemoglobin [43], d) an enzyme (MnP) that is degraded by its own substrate [45], e) an ion channel (CFTR) that adopts multiple conformations [15], and f) oxygen consumption in a living tissue (fish brain) [21]. The first three systems were anticipated to be homogeneous, and our method confirmed this. Whereas, the final three systems were suspected to be heterogeneous. Our results suggested that the MnP and CFTR systems were indeed heterogeneous, and their non-MM-like curves could not be explained by cooperativity between binding sites. The case of the living tissue is more complex and, although the standard MM model was grossly inadequate, we could not unambiguously resolve whether negative cooperation or heterogeneity made a better explanation. The heterogeneous Michaelis-Menten model is explanatory, however its approximation of heterogeneity is not based on any first principles (aside from the general observation that many things in nature are approximately normally distributed). In many cases, a specialised model would be preferred, such as a multi-state model for ion channels [15], gene regulation [50], or transcription elongation [51].

Historically, scientists have taken great strides to actively avoid non-homogeneous enzyme preparations, so as to eliminate as much background noise as possible. However, in some cases heterogeneous preparations may be preferable. For example one may wish to purify a large sample of computationally modelled enzyme sequences, such as ancestral reconstructions, and screen them simultaneously for catalytic activity. Moreover, capturing variability may be essential for characterising primordial proteins, which may have existed as populations of “quasi-species” [52] produced by an ambiguous genetic code [10]. Our results here have paved the way for these kinds of experiments. Heterogeneity is everywhere in nature and identifying it is only the first challenge. The greater challenge lies in harnessing heterogeneity experimentally and determining when it is biologically meaningful [20].

## Data Availability

Our source code is available at https://github.com/jordandouglas/HetMM.

## Acknowledgements

This work was funded by the Alfred P. Sloan Foundation Matter-to-Life program Grant number G-2021-16944. The authors acknowledge the use of New Zealand eScience Infrastructure (NeSI) high performance computing facilities, funded jointly by NeSI’s collaborator institutions and the Ministry of Business, Innovation & Employment’s Research Infrastructure programme. The authors thank Jules Devaux and Tony Hickey for sharing their data on oxygen consumption in fish.

## Supporting Information

**Fig S1.**
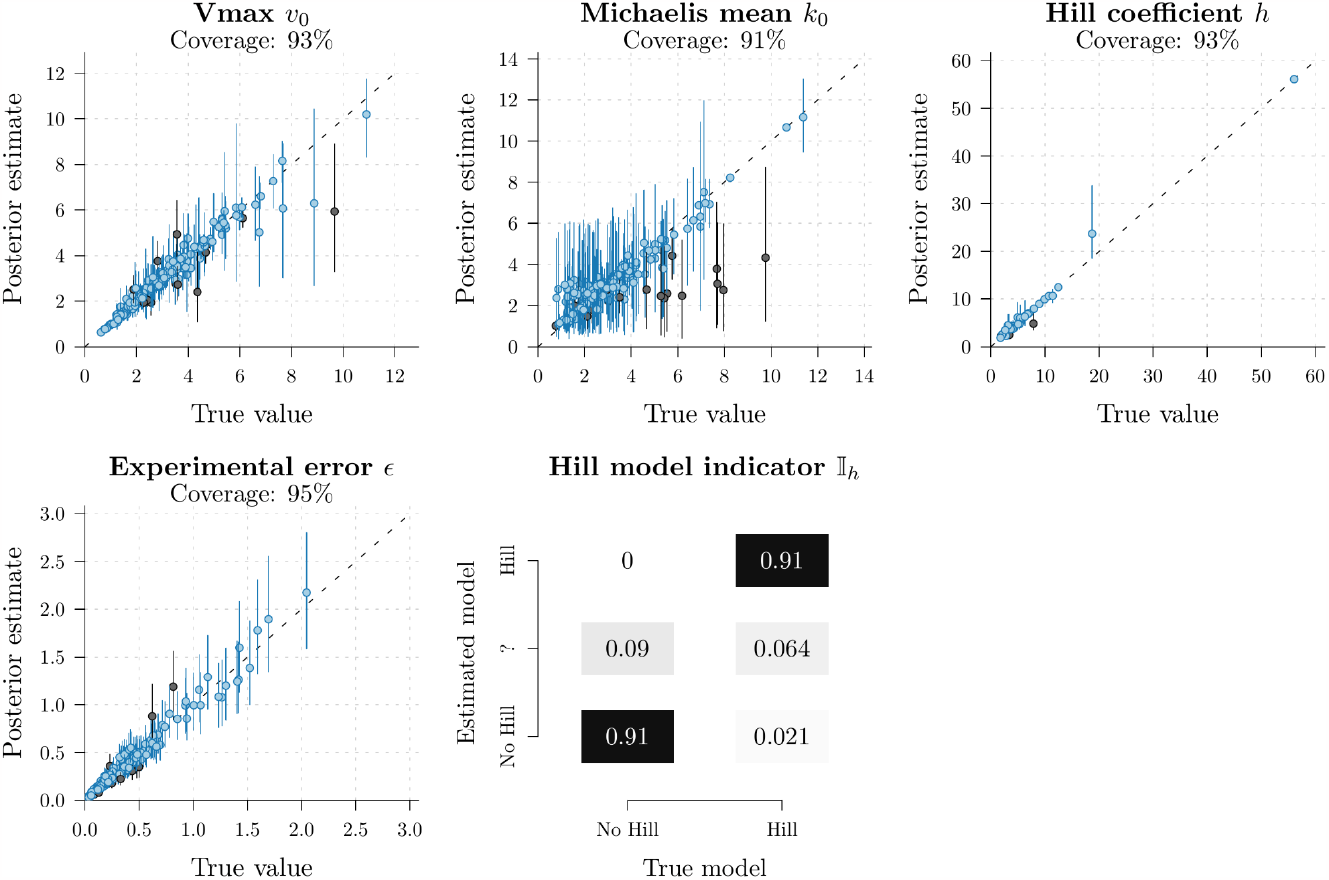
Well-calibrated simulation study on the homogeneous Hill model. Each point represents an MCMC chain performed on one of two hundred datasets simulated under known parameters. h is part of the Hill model, and therefore its coverage is only calculated when p(Ih≠0) > 0.95. The true estimated model is indicated if it has more than 0.95 posterior support, while ‘?’ denotes uncertainty (i.e., 0.05 < p(Ih≠0) < 0.95).

## References

[1] L. Michaelis, M. L. Menten, and others, “Die kinetik der invertinwirkung,” Biochem. z, vol. 49, nos. 333-369, p. 352, 1913.

[2] B. Srinivasan, “A guide to the michaelis–menten equation: Steady state and beyond,” The FEBS journal, vol. 289, no. 20, pp. 6086–6098, 2022.

[3] S. Schnell, “Validity of the michaelis–menten equation–steady-state or reactant stationary assumption: That is the question,” The FEBS journal, vol. 281, no. 2, pp. 464–472, 2014.

[4] G. E. Briggs and J. B. S. Haldane, “A note on the kinetics of enzyme action,” Biochemical journal, vol. 19, no. 2, p. 338, 1925.

[5] W. A. Susor, M. Kochman, and W. J. Rutter, “Heterogeneity of presumably homogeneous protein preparations,” Science, vol. 165, no. 3899, pp. 1260–1262, 1969.

[6] S. Brown, N. Muhamad, K. C. Pedley, and D. C. Simcock, “The kinetics of enzyme mixtures,” Molecular Biology Research Communications, vol. 3, no. 1, p. 21, 2014.

[7] N. Krahn, D. Söll, and O. Vargas-Rodriguez, “Diversification of aminoacyl-tRNA synthetase activities via genomic duplication,” Frontiers in Physiology, vol. 13, p. 1704, 2022.

[8] B. Errede and M. D. Kamen, “Comparative kinetic studies of cytochromes c in reactions with mitochondrial cytochrome c oxidase and reductase,” Biochemistry, vol. 17, no. 6, pp. 1015–1027, 1978.

[9] M. H. Schwartz and T. Pan, “Temperature dependent mistranslation in a hyperthermophile adapts proteins to lower temperatures,” Nucleic acids research, vol. 44, no. 1, pp. 294–303, 2015.

[10] C. W. Carter Jr and P. R. Wills, “The roots of genetic coding in aminoacyl-tRNA synthetase duality,” Annual review of biochemistry, vol. 90, pp. 349–373, 2021.

[11] J. Douglas, R. Bouckaert, C. Carter, and P. R. Wills, “Enzymic recognition of amino acids drove the evolution of primordial genetic codes,” Research Square, 2023.

[12] H. Frauenfelder, S. G. Sligar, and P. G. Wolynes, “The energy landscapes and motions of proteins,” Science, vol. 254, no. 5038, pp. 1598–1603, 1991.

[13] S. W. Englander and L. Mayne, “The nature of protein folding pathways,” Proceedings of the National Academy of Sciences, vol. 111, no. 45, pp. 15873–15880, 2014.

[14] Y. Luo et al., “Resolving molecular heterogeneity with single-molecule centrifugation,” Journal of the American Chemical Society, vol. 145, no. 6, pp. 3276–3282, 2023.

[15] F. Liu, Z. Zhang, L. Csanády, D. C. Gadsby, and J. Chen, “Molecular structure of the human CFTR ion channel,” Cell, vol. 169, no. 1, pp. 85–95, 2017.

[16] K. Vamvaca, B. Vögeli, P. Kast, K. Pervushin, and D. Hilvert, “An enzymatic molten globule: Efficient coupling of folding and catalysis,” Proceedings of the National Academy of Sciences, vol. 101, no. 35, pp. 12860–12864, 2004.

[17] H. S. Tehrani and A. A. Moosavi-Movahedi, “Catalase and its mysteries,” Progress in Biophysics and Molecular Biology, vol. 140, pp. 5–12, 2018.

[18] M. J. Davies, “Protein oxidation and peroxidation,” Biochemical journal, vol. 473, no. 7, pp. 805–825, 2016.

[19] S. Ahmad et al., “Protein oxidation: An overview of metabolism of sulphur containing amino acid, cysteine,” Frontiers in Bioscience-Scholar, vol. 9, no. 1, pp. 71–87, 2017.

[20] S. J. Altschuler and L. F. Wu, “Cellular heterogeneity: Do differences make a difference?” Cell, vol. 141, no. 4, pp. 559–563, 2010.

[21] J. B. Devaux, C. P. Hedges, N. Birch, N. Herbert, G. M. Renshaw, and A. J. Hickey, “Electron transfer and ros production in brain mitochondria of intertidal and subtidal triplefin fish (tripterygiidae),” Journal of Comparative Physiology B, pp. 1–12, 2023.

[22] D. P. Anderson et al., “Evolution of an ancient protein function involved in organized multicellularity in animals,” Elife, vol. 5, p. e10147, 2016.

[23] R. N. Randall, C. E. Radford, K. A. Roof, D. K. Natarajan, and E. A. Gaucher, “An experimental phylogeny to benchmark ancestral sequence reconstruction,” Nature communications, vol. 7, no. 1, p. 12847, 2016.

[24] J. J. Hobson, Z. Li, H. Hu, and C. W. Carter Jr, “A Leucyl-tRNA Synthetase Urzyme: Authenticity of tRNA Synthetase Catalytic Activities and Promiscuous Phosphorylation of Leucyl-5’ AMP,” International Journal of Molecular Sciences, vol. 23, no. 8, p. 4229, 2022.

[25] L. Li, V. Weinreb, C. Francklyn, and C. W. Carter, “Histidyl-tRNA synthetase urzymes: Class I and II aminoacyl tRNA synthetase urzymes have comparable catalytic activities for cognate amino acid activation,” Journal of Biological Chemistry, vol. 286, no. 12, pp. 10387–10395, 2011.

[26] A. F. Kolodziej, T. Tan, and D. E. Koshland, “Producing positive, negative, and no cooperativity by mutations at a single residue located at the subunit interface in the aspartate receptor of salmonella typhimurium,” Biochemistry, vol. 35, no. 47, pp. 14782–14792, 1996.

[27] S. Hinderlich, R. Stasche, R. Zeitler, and W. Reutter, “A bifunctional enzyme catalyzes the first two steps in n-acetylneuraminic acid biosynthesis of rat liver,” Journal of Biological Chemistry, vol. 272, no. 39, pp. 24313–24318, 1997.

[28] A. V. Hill, “The possible effects of the aggregation of the molecules of hemoglobin on its dissociation curves,” j. physiol., vol. 40, pp. iv–vii, 1910.

[29] J. N. Weiss, “The hill equation revisited: Uses and misuses,” The FASEB Journal, vol. 11, no. 11, pp. 835–841, 1997.

[30] C. Y. Seow, “Hill’s equation of muscle performance and its hidden insight on molecular mechanisms,” Journal of General Physiology, vol. 142, no. 6, pp. 561–573, 2013.

[31] H. Abeliovich, “An empirical extremum principle for the hill coefficient in ligand-protein interactions showing negative cooperativity,” Biophysical journal, vol. 89, no. 1, pp. 76–79, 2005.

[32] R. Bouckaert et al., “BEAST 2.5: An advanced software platform for Bayesian evolutionary analysis,” PLoS computational biology, vol. 15, no. 4, p. e1006650, 2019.

[33] Z. Yang and C. E. Rodríguez, “Searching for efficient markov chain monte carlo proposal kernels,” Proceedings of the National Academy of Sciences, vol. 110, no. 48, pp. 19307–19312, 2013.

[34] G. Baele, P. Lemey, A. Rambaut, and M. A. Suchard, “Adaptive mcmc in bayesian phylogenetics: An application to analyzing partitioned data in beast,” Bioinformatics, vol. 33, no. 12, pp. 1798–1805, 2017.

[35] J. Douglas, R. Zhang, and R. Bouckaert, “Adaptive dating and fast proposals: Revisiting the phylogenetic relaxed clock model,” PLoS computational biology, vol. 17, no. 2, p. e1008322, 2021.

[36] A. Rambaut, A. J. Drummond, D. Xie, G. Baele, and M. A. Suchard, “Posterior summarization in Bayesian phylogenetics using tracer 1.7,” Systematic biology, vol. 67, no. 5, p. 901, 2018.

[37] S. Talts, M. Betancourt, D. Simpson, A. Vehtari, and A. Gelman, “Validating bayesian inference algorithms with simulation-based calibration,” arXiv preprint arXiv:1804.06788, 2018.

[38] S. T. Radev et al., “OutbreakFlow: Model-based bayesian inference of disease outbreak dynamics with invertible neural networks and its application to the covid-19 pandemics in germany,” PLoS computational biology, vol. 17, no. 10, p. e1009472, 2021.

[39] J. Douglas, C. L. Jiménez-Silva, and R. Bouckaert, “StarBeast3: Adaptive Parallelized Bayesian Inference under the Multispecies Coalescent,” Systematic Biology, vol. 71, no. 4, pp. 901–916, 2022.

[40] V. Jiménez-Jiménez, C. Martí-Gómez, M. Á. del Pozo, E. Lara-Pezzi, and F. Sánchez-Cabo, “Bayesian inference of gene expression,” Exon Publications, pp. 65–87, 2021.

[41] K. A. Johnson and R. S. Goody, “The original michaelis constant: Translation of the 1913 michaelis–menten paper,” Biochemistry, vol. 50, no. 39, pp. 8264–8269, 2011.

[42] J. C. Aledo, “Renz: An r package for the analysis of enzyme kinetic data,” BMC bioinformatics, vol. 23, no. 1, p. 182, 2022.

[43] J. W. Severinghaus, “Simple, accurate equations for human blood o2 dissociation computations,” Journal of Applied Physiology, vol. 46, no. 3, pp. 599–602, 1979.

[44] C. Anstey, “A new model for the oxyhaemoglobin dissociation curve,” Anaesthesia and intensive care, vol. 30, no. 5, pp. 376–387, 2002.

[45] C. Palma, M. Moreira, G. Feijoo, and J. Lema, “Enhanced catalytic properties of mnp by exogenous addition of manganese and hydrogen peroxide,” Biotechnology letters, vol. 19, pp. 263–268, 1997.

[46] M. Sundaramoorthy, M. H. Gold, and T. L. Poulos, “Ultrahigh (0.93 Å) resolution structure of manganese peroxidase from phanerochaete chrysosporium: Implications for the catalytic mechanism,” Journal of inorganic biochemistry, vol. 104, no. 6, pp. 683–690, 2010.

[47] K. L. Gunderson and R. R. Kopito, “Conformational states of cftr associated with channel gating: The role of atp binding and hydrolysis,” Cell, vol. 82, no. 2, pp. 231–239, 1995.

[48] D. C. Gadsby, P. Vergani, and L. Csanády, “The abc protein turned chloride channel whose failure causes cystic fibrosis,” Nature, vol. 440, no. 7083, pp. 477–483, 2006.

[49] J. C. Cheung, P. K. Chiaw, S. Pasyk, and C. E. Bear, “Molecular basis for the atpase activity of cftr,” Archives of biochemistry and biophysics, vol. 476, no. 1, pp. 95–100, 2008.

[50] D. Chu, N. R. Zabet, and B. Mitavskiy, “Models of transcription factor binding: Sensitivity of activation functions to model assumptions,” Journal of Theoretical Biology, vol. 257, no. 3, pp. 419–429, 2009.

[51] J. Douglas, R. Kingston, and A. J. Drummond, “Bayesian inference and comparison of stochastic transcription elongation models,” PLoS computational biology, vol. 16, no. 2, p. e1006717, 2020.

[52] P. R. Wills, K. Nieselt, and J. S. McCaskill, “Emergence of coding and its specificity as a physico-informatic problem,” Origins of Life and Evolution of Biospheres, vol. 45, pp. 249–255, 2015.

